# Local Ancestry Prediction with *PyLAE*

**DOI:** 10.1101/2020.11.13.380105

**Authors:** Alexander Smetanin, Nikita Moshkov, Tatiana V. Tatarinova

## Abstract

**Summary:** We developed PyLAE - a new tool for determining local ancestry along a genome using whole-genome sequencing data or high-density genotyping experiments. PyLAE can process an arbitrarily large number of ancestral populations (with or without an informative prior). Since PyLAE does not involve estimation of many parameters, it can process thousands of genomes within a day. Computational efficiency, straightforward presentation of results, and an ease of installation makes *PyLAE* a useful tool to study admixed populations.

**Availability and implementation:** The source code and installation manual are available at https://github.com/smetam/pylae.

## Introduction

In association studies, researchers combine samples with different (usually, opposing) phenotypes and compare Single Nucleotide Polymorphisms (SNPs) frequencies in two groups. When enough samples are available (typically, thousands for complex traits), a significant difference in frequencies between the groups suggests an association between the position on the genome and the studied phenotype. However, there is a possibility that the association is due to inhomogeneity of the study group in terms of provenance/origin (for example, all people with the disease are of French origin, and the healthy cohort is Bulgarian). In this case, two populations may have different frequencies of ancestry informative markers (AIM) that are not causal to the phenotype.

An intuitive approach to solving this problem is determining the population structure first and then adjusting for it. However, this strategy is complicated for admixed populations. Due to meiotic recombination during transmission of genetic material to the offspring, individual segments of the genome may have different origins. Recombination makes it hard to determine the source of such individuals (or plant populations), especially locus-specific local origin.

There are several existing solutions to this problem, such as (LAMP, LAMP-ANC, and RFMIX). However, there is always room to develop a user-friendly, fast, accurate approach. LAMP [1] algorithm determines local origin in mixed populations. The LAMP input includes recombination rate, global mixing ratio, and the number of generations that have passed since the mixing started. The recombination rate can be considered known due to previous studies [2], while the global proportion can be estimated using, for example, the ADMIXTURE tool [3]. The idea of the LAMP method is as follows. The iterated conditional modes clustering algorithm (ICM) determines the likelihood that a segment of the genome within a selected window has a specific origin. An individual SNP is assigned the source by the “popular vote” approach using the inferred origins of all windows covering this position. ICM is a modification of the Expectation-Maximization (EM) algorithm, which receives a point estimate at the E-step under the assumption that the a priori estimates are sufficiently accurate. Therefore, the ICM is faster than the EM approach. To construct an accurate *a priori* estimate, LAMP uses the MAXVAR algorithm, which works only for two populations. The size of the genomic window is selected to minimize classification errors. LAMP works fast, but its accuracy decreases if the ethnic admixture ratio is close to 1:1.

LAMP-ANC [4] is a modification of the LAMP tool, showing a higher accuracy than LAMP. This modification also allows triple mixing to be estimated, while LAMP cannot determine frequencies for more than two ancestral populations.

Machine Learning algorithm RFMIX [5, 6] treats origin as a hidden parameter in its statistical model. RFMIX employs conditional random fields and decision trees. The RFMIX model does not impose restrictions on the number of ancestral populations and types of mixing.

We developed PyLAE - a new tool for determining local origin along a genome using whole-genome sequencing data or high-density genotyping experiments. PyLAE can process an arbitrarily large number of ancestral populations (with or without an informative prior). PyLAE does not involve the estimation of many parameters. *PyLAE* is a useful tool to study admixed populations [7–9]. We have tested this approach using the 1000 Genomes database.

## Materials and Methods

### Data source

Whole-genome sequencing data in the VCF format were obtained from the 1000 genomes project (ftp://ftp.1000genomes.ebi.ac.uk/vol1/ftp/, genome version GRCh37). We used the dataset of 2504 worldwide individuals with the reported ethnic origin [10] and phased genome sequence data.

### Data preprocessing

#### Admixture

To confirm the reported origin, we have extracted 130,000 ancestry informative markers identified by a worldwide study contacted by the National Genographic consortium, as described in [11]. The supervised admixture analysis was performed using the K=9 component division of ancestral populations into the following categories: North-East Asian, Mediterranean, South African, South-West Asian, Native American, Oceanian, Southeast Asian, Northern European, and Sub-Saharan African components. We used the putative ancestral populations from our earlier study [11]. The admixture components represent proportions of an individual’s genotype attributed to each of the nine putative ancestral genomes. The obtained nine-dimensional vectors were clustered based on the L2 norm (Euclidean distance). The optimal number of clusters was determined using the weighted Kullback-Leibler distance approach [12, 13]. Within each group, the admixture profiles of individuals are similar. This dimension-reduction step allows the identification of potentially admixed individuals. This assignment was validated using haplogroup information and reAdmix analysis in group mode [14]. reAdmix algorithm represents an individual as a weighted sum of present-day populations (e.g., 50% British, 25% Russian, 25% Han Chinese) based on K admixture components. Instead of attempting to solve an “exact admixture” problem, we aim to find the smallest subset of populations whose combined admixture components are close to those of the individuals within a small tolerance margin. Due to the range of natural variation, the admixture proportions can be considered exact neither for the reference populations nor for the tested individuals. The admixture proportions we use are merely maximum likelihood estimates and may fail to be exactly equal to the actual shares of ancestral genomes. Therefore, determining ethnic proportions is mathematically and computationally more challenging than finding a single most fitting bio-origin. From our experience with reAdmix, this tool accurately identifies all significant components; however, small proportions are frequently mis-assigned. When similar individuals are analyzed as a group, it is possible to estimate even the small proportions reliably.

#### Phasing

The phased data were obtained for the 1000 genomes from the open repository ftp://ftp.1000genomes.ebi.ac.uk/vol1/ftp/. Sudmant et al. [10] used short-read Illumina DNA sequencing data and statistically phased onto haplotype blocks in 26 human populations.

#### Two modes

We have developed two modes: ‘normal’ and ‘haploid.’ For the ‘normal’ mode, we used the samples as-is. For the ‘haploid’ mode, we used the phasing information provided by the 1000 Genomes and assume that the parents were homozygous. Thus, we generated two homozygous parents from each sample.

### Local Ancestry Bayesian Approach (*PyLAE*)

#### Input

**Individual**: VCF file and (optional) Admixture vector obtained by supervised Admixture, as described above.
**Reference**: multi VCF file with putative ancestral populations corresponding to K component admixture. N is the number of positions overlapping between the reference and the individual.

The number of genomic positions should be significantly large to ensure dense genomic coverage, while not being too large, so the distance between two consecutive points is above the LD for the studied organism.

#### Stage 1. Bayesian posterior probability

Posterior probability *P*(*population*|*genotype*)is calculated using the Bayes formula:

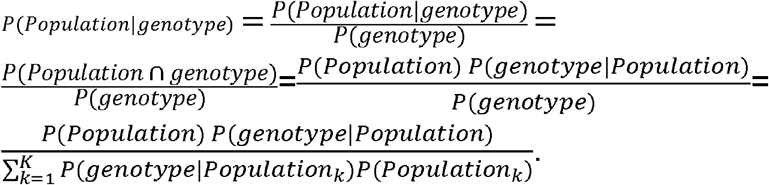

The prior probability of a population *P*(*population*) is equal to the analyzed individual’s admixture vector. We assume that adjacent positions have the same origin and the origin of all positions within a window of length ***L***. Since the distance between two consecutive ancestry informative markers is above LD, positions within the same window are independent.

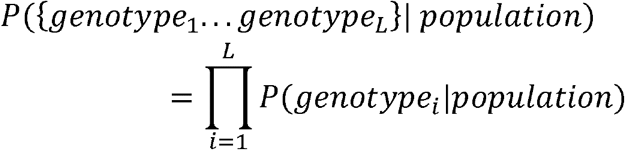

Therefore

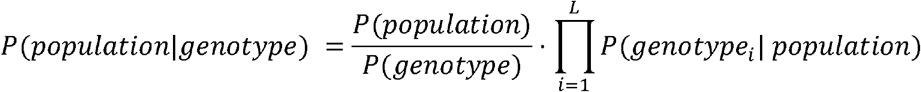

*P*(*genotype_i_|population*) is estimated from observations, using genotypes of putative ancestral populations in position *i*

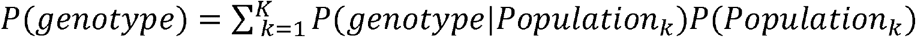

Therefore, we have K posterior probabilities for every genotype, and the matrix of posterior probabilities *T* × *K*, where *T* = ^*N*^/*L*.

We are interested in determining the optimal sequence of the source populations along the genome. This problem is solved using the Viterbi algorithm [15]. For computational efficiency, all calculations are performed in the log-space.

#### Stage 2. Viterbi algorithm

Our state-space *S* consists of *K* ancestral populations. Population labels (*i*=1..*K*) are the hidden states in our model. The initial probabilities *π_i_* of being in the i^th^ hidden state can either be assumed to be uniform with 1/K probability of each population or set to be equal to global admixture components. Transition probabilities *α_i,j_* of transitioning from the state *i* to *j* are inversely proportional to the TreeMix distances between corresponding ancestral populations. Transition probability from state *i* to *j a_ij_* can be calculated from distances between ancestral populations. The emission probabilities *b_j_*(*O*) is calculated by the Bayes formula (above). At the first step of the algorithm *δ_j_*(1) = *π_j_b_j_*(*O*_1_). At each next step *δ_j_*(*t* + 1) = *max_i_ δ_i_*(*t*) *a_ij_*(*O*_*t*+1_). The pointers *D_j_*(*t* + 1) are stored for tracing back. *D_j_*(*t* + 1) = *argmax_i_ δ_i_*(*t*) *a_ij_*(*O*_*t* + 1_). At the terminal state: *S_T_* = *argmax_i_ δ_i_*(*T*), *S_t_* = *D*_*S*_*t*+1__(*t* + 1).

For efficiency, the computation is performed in the log-space. The algorithm is implemented as a python script and can be adapted to analyze any organism using user-provided ancestral components, prior probabilities, and transition matrix.

### Application of Local Ancestry

Most of the currently living individuals are admixed to various degrees. To compare genotypes between populations and analyze signals of selection, it is essential to identify and exclude introgressed regions containing non-representative genotypes. We have determined local ancestry profiles for all 2504 analyzed individuals from the 1000 Genomes project. With our approach, even the admixed individuals that are typically excluded from the analysis (therefore, reducing the statistical power of the study) can be retained after masking the introgressed regions.

We have used the approach of Chekalin et al. [16–18] and annotated the VCFs with the ANNOVAR tool using the hg19 human genome annotation and the refGene database (http://varianttools.sourceforge.net/Annotation/RefGene). The SNPs were classified by their location and function: 58% intergenic, 38% intronic, 5% ncRNA, and 1% exonic (64% of them are synonymous, and 36% are nonsynonymous). Next, we counted the synonymous and nonsynonymous SNPs per each KEGG pathway [16, 19–21].

It is reasonable to assume that pathways accumulate nonsynonymous SNPs at the same rate during neutral evolution; therefore, the pathways’ enrichment scores can be approximated by a normal distribution. The pathways under selection will appear as outliers.

To analyze differences in numbers of SNPs per pathway between the two groups of populations (groups A and B), we need to calculate: 1) DSSE scores (synonymous) and 2) DNSE scores (nonsynonymous) [16]. Therefore, we perform the following steps:

1. Find the number of (non)synonymous SNPs in groups A and B.

a. Let *I* represent the total number of studied pathways, and *i*□=□1,…,*I*, be the number of the (non)synonymous SNP per i^th^ pathway are nS(i) and nA(i). The expected fraction of (non)synonymous SNPs in Siberian individuals is given by 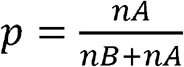, where *nA* is the amount of A (non)synonymous SNPs in all KEGG pathways, *nB* is the amount of B (non) synonymous SNPs in all KEGG pathways. The fraction p_i_ of A (non)synonymous SNPs in the i^th^ KEGG pathway is 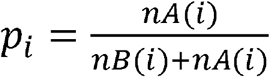.
2. The enrichment D(N)SE scores are computed for every pathway with continuity correction:

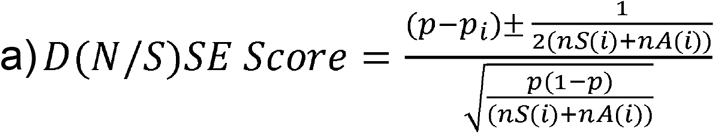
3. P-values are calculated using Bonferroni and Benjamini–Hochberg corrections and identified the differentially enriched pathways. A pathway is differentially enriched if the absolute value of the D(N/S)SE score >4, and the adjusted P-value <0.005 [22]. A pathway is differentially enriched if the absolute value of the D(N/S)SE score >4, and the adjusted P-value <0.005.
4. To consider the excess of synonymous SNPs over nonsynonymous SNPs, we calculate enrichment scores for synonymous SNPs, DSSE. Bonferroni-adjusted P-values are log-transformed and multiplied by the sign of the D(N/S)SE statistic so that positive scores correspond to enrichment in group B. Negative scores correspond to enrichment in group A. As a condition of significance, and we require the following: The P-value of the nonsynonymous test was below the P-value of the synonymous test for each pathway. Besides, it was needed for the Bonferroni-corrected nonsynonymous test P-value to be below 0.01. For each pathway, the synonymous test’s P-value was above the P-value of the corresponding nonsynonymous test.

## Results

### 1. Investigation of the reference dataset

Nine-dimensional admixture profiles were calculated for 2,504 individuals (130K markers each) using supervised Admixture. Every sample was analyzed first in diploid and then in haploid mode (Figure 1).

**Figure 1:**
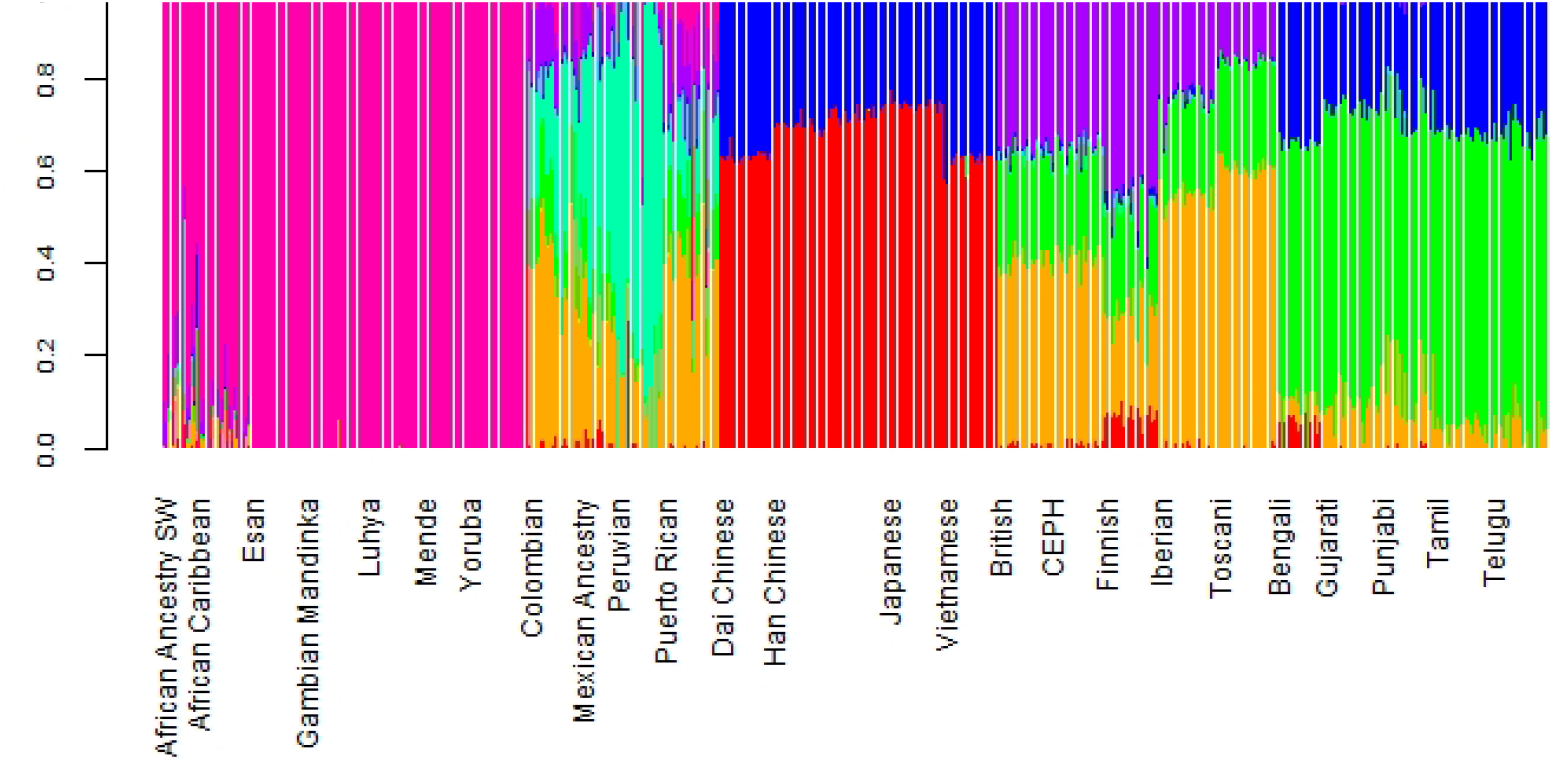
Admixture for 10000 Genomes individuals. Top: diploid mode. Bottom: haploid mode. Colors: Red - North-East Asian, Brown - Mediterranean, Light green - South African, Dark green - South-West Asian, Light blue - Native American, Blue - Oceanian, Dark blue - Southeast Asian, Purple - Northern European, Magenta - Sub-Saharan African.

Diploid and haploid admixture profiles were clustered within each reported ethnic origin (Figure 2, Tables 1 and 2). African Ancestry SW, African Caribbean, Columbian, Gambian Mandinka, Mexican Ancestry, Peruvian, and Puerto-Rican showed 2-3 subgroups within each reported ethnicity, which is consistent with the complex population history of these populations.

**Figure 2:**
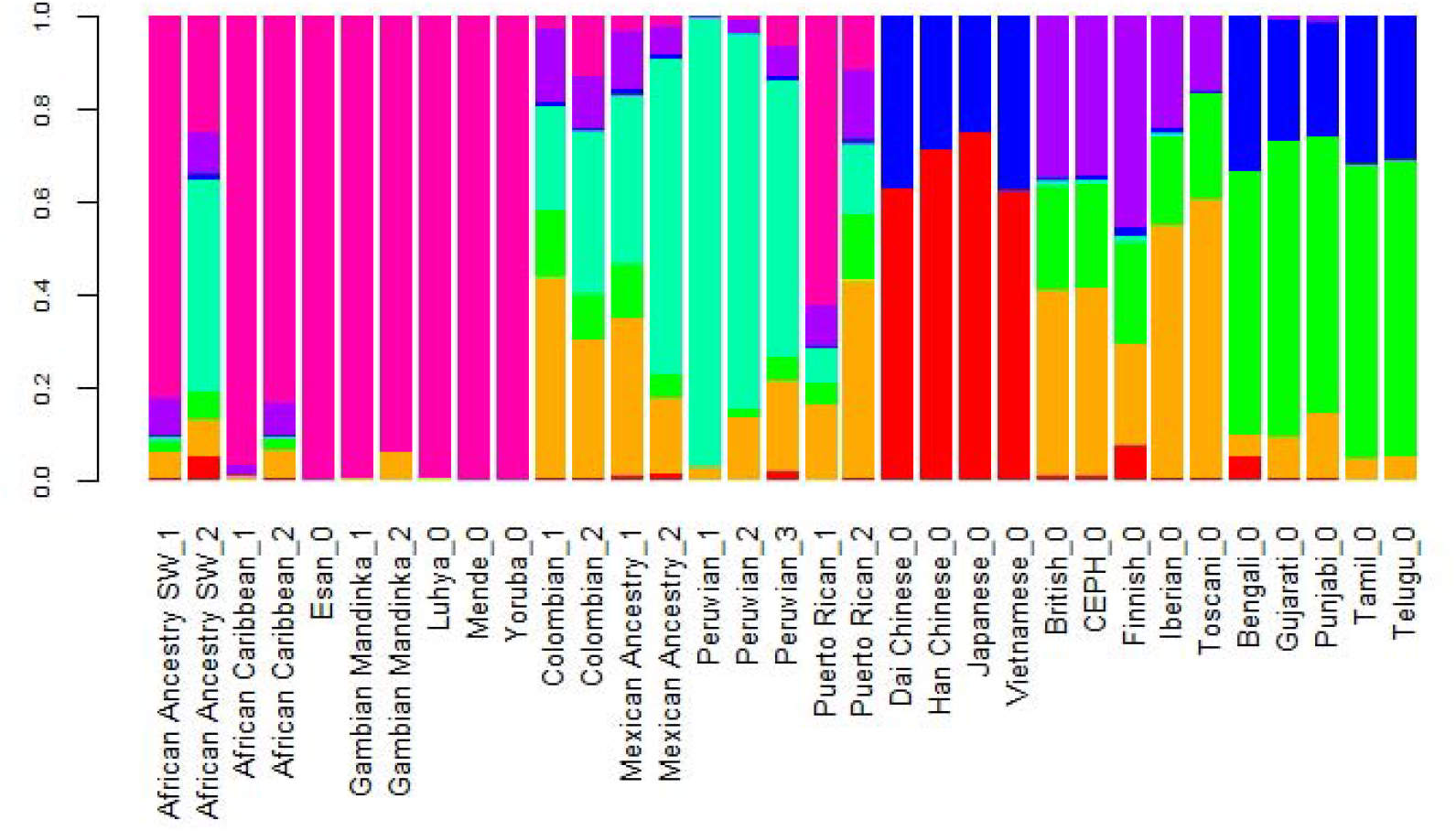
Clustered Admixture for 10000 Genomes individuals. Top: diploid mode. Bottom: haploid mode. Colors: Red - North-East Asian, Brown - Mediterranean, Light green - South African, Dark green - South-West Asian, Light blue - Native American, Blue - Oceanian, Dark blue - Southeast Asian, Purple - Northern European, Magenta - Sub-Saharan African.

**Table 1.**
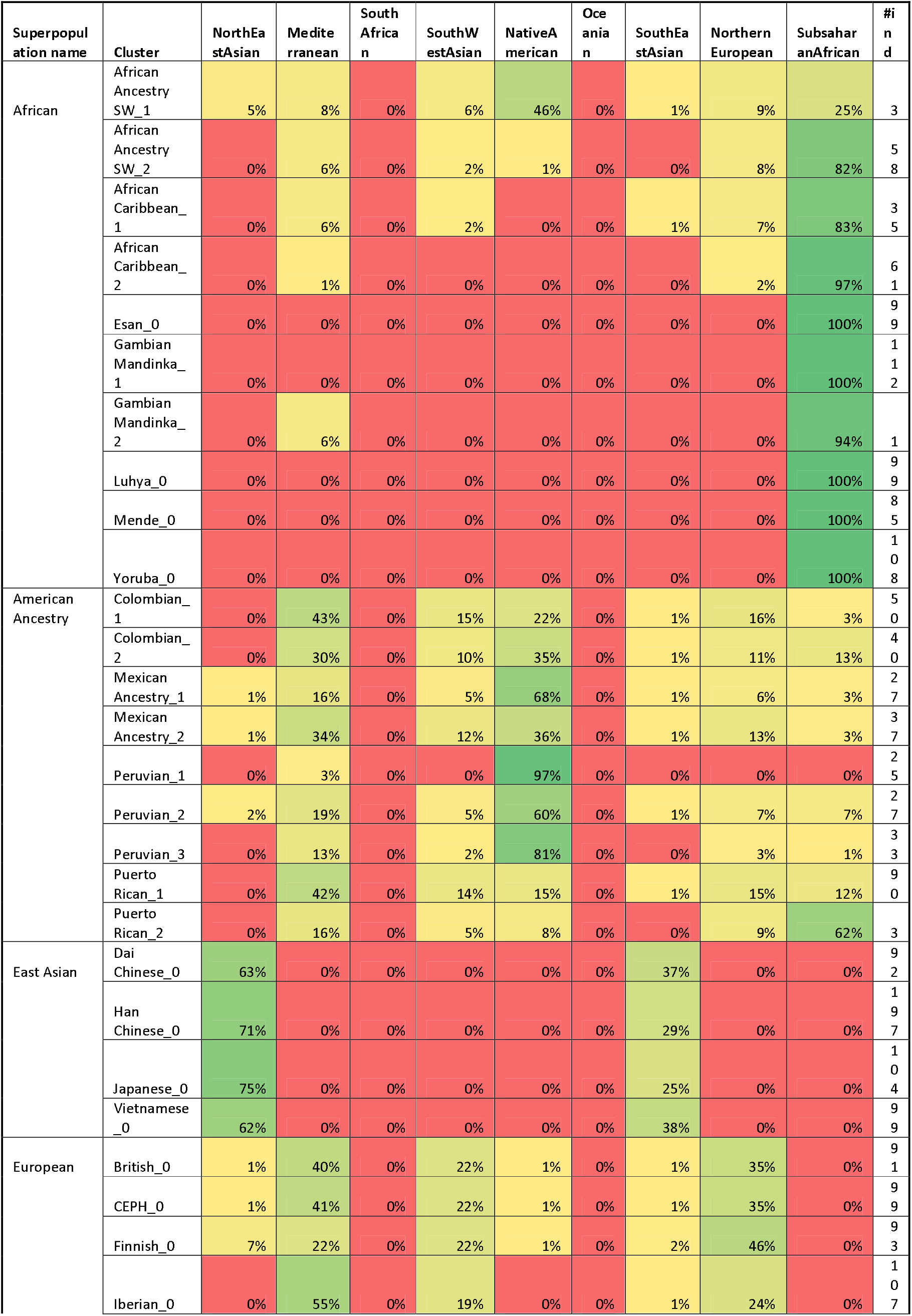
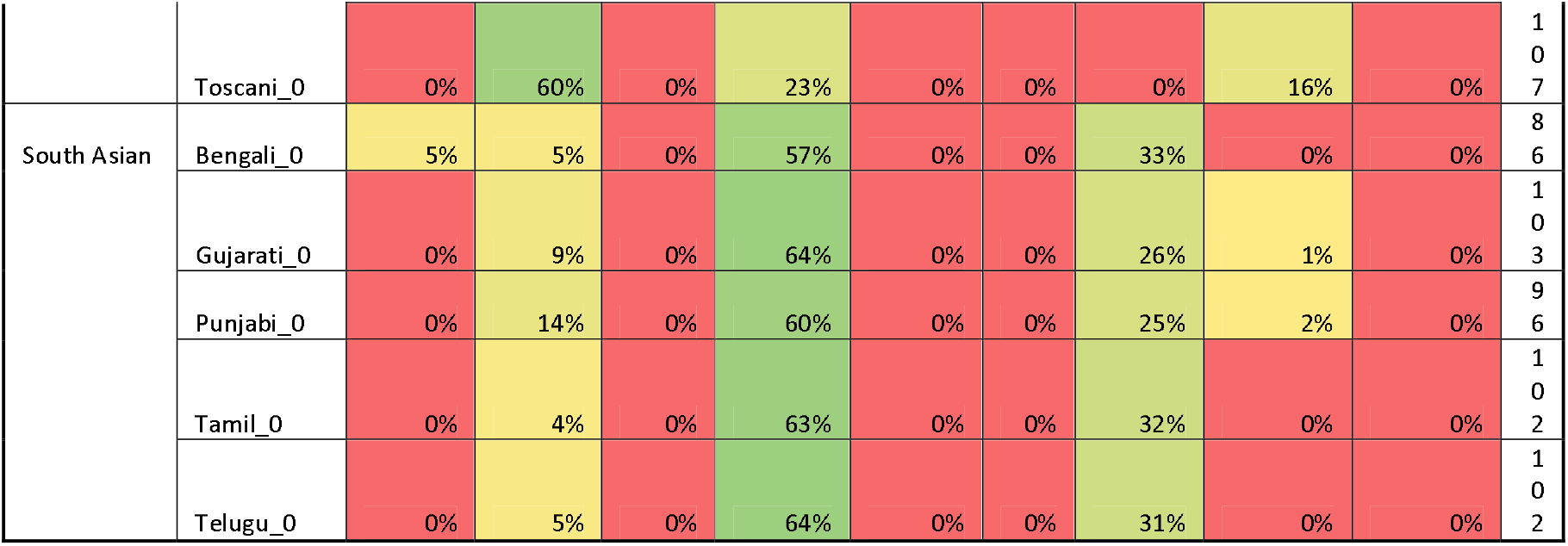
Admixture vector clusters for 1000 Genomes individuals

**Table 2.** Haploid admixture vector clusters for 1000 Genomes individuals

| Superpopulation name | Cluster | NorthEastAsian | Mediterranean | South African | SouthWestAsian | Native American | Oceania | SouthEastAsian | Northern European | Subsaharan African | #parents |
| --- | --- | --- | --- | --- | --- | --- | --- | --- | --- | --- | --- |
| African | African Ancestry SW_1 | 0% | 0% | 0% | 0% | 0% | 0% | 0% | 2% | 98% | 71 |
|  | African Ancestry SW_2 | 0% | 4% | 0% | 1% | 1% | 0% | 0% | 15% | 78% | 45 |
|  | African Ancestry SW_3 | 0% | 9% | 0% | 2% | 54% | 0% | 0% | 8% | 27% | 6 |
|  | African Caribbean_1 | 0% | 14% | 0% | 10% | 0% | 0% | 5% | 10% | 62% | 6 |
|  | African Caribbean_2 | 0% | 1% | 0% | 0% | 0% | 0% | 0% | 8% | 90% | 26 |
|  | African Caribbean_3 | 0% | 0% | 0% | 0% | 0% | 0% | 0% | 0% | 100% | 160 |
|  | Esan_0 | 0% | 0% | 0% | 0% | 0% | 0% | 0% | 0% | 100% | 198 |
|  | Gambian Mandinka_0 | 0% | 0% | 0% | 0% | 0% | 0% | 0% | 0% | 100% | 226 |
|  | Luhya_0 | 0% | 0% | 0% | 0% | 0% | 0% | 0% | 0% | 100% | 198 |
|  | Mende_0 | 0% | 0% | 0% | 0% | 0% | 0% | 0% | 0% | 100% | 170 |
|  | Yoruba_0 | 0% | 0% | 0% | 0% | 0% | 0% | 0% | 0% | 100% | 216 |
| American | Colombian_1 | 0% | 43% | 0% | 6% | 38% | 0% | 0% | 2% | 11% | 86 |
|  | Colombian_2 | 0% | 60% | 0% | 12% | 22% | 0% | 0% | 4% | 1% | 94 |
|  | Mexican Ancestry_1 | 0% | 35% | 0% | 3% | 59% | 0% | 0% | 1% | 2% | 60 |
|  | Mexican Ancestry_2 | 0% | 53% | 0% | 10% | 33% | 0% | 0% | 4% | 1% | 46 |
|  | Mexican Ancestry_3 | 0% | 9% | 0% | 0% | 91% | 0% | 0% | 0% | 0% | 22 |
|  | Peruvian_1 | 1% | 27% | 0% | 1% | 64% | 0% | 1% | 1% | 6% | 48 |
|  | Peruvian_2 | 0% | 14% | 0% | 0% | 84% | 0% | 0% | 0% | 1% | 50 |
|  | Peruvian_3 | 0% | 0% | 0% | 0% | 100% | 0% | 0% | 0% | 0% | 72 |
|  | Puerto Rican_1 | 0% | 59% | 0% | 12% | 16% | 0% | 0% | 3% | 10% | 180 |
|  | Puerto Rican_2 | 0% | 16% | 0% | 1% | 6% | 0% | 0% | 9% | 67% | 6 |
| East Asian | Dai Chinese_0 | 66% | 0% | 0% | 0% | 0% | 0% | 34% | 0% | 0% | 184 |
|  | Han Chinese_0 | 77% | 0% | 0% | 0% | 0% | 0% | 23% | 0% | 0% | 394 |
|  | Japanese_0 | 82% | 0% | 0% | 0% | 0% | 0% | 18% | 0% | 0% | 208 |
|  | Vietnamese_0 | 65% | 0% | 0% | 0% | 0% | 0% | 35% | 0% | 0% | 198 |
| European | British_0 | 0% | 46% | 0% | 21% | 0% | 0% | 0% | 34% | 0% | 182 |
|  | CEPH_0 | 0% | 47% | 0% | 20% | 0% | 0% | 0% | 33% | 0% | 198 |
|  | Finnish_0 | 7% | 17% | 0% | 22% | 0% | 0% | 1% | 53% | 0% | 186 |
|  | Iberian_0 | 0% | 72% | 0% | 16% | 0% | 0% | 0% | 12% | 0% | 214 |
|  | Toscani_0 | 0% | 82% | 0% | 18% | 0% | 0% | 0% | 0% | 0% | 214 |
| South Asian | Bengali_0 | 0% | 0% | 0% | 64% | 0% | 0% | 36% | 0% | 0% | 172 |
|  | Gujarati_1 | 0% | 0% | 0% | 74% | 0% | 0% | 26% | 0% | 0% | 180 |
|  | Gujarati_2 | 0% | 12% | 0% | 70% | 0% | 0% | 18% | 0% | 0% | 26 |
|  | Punjabi_1 | 0% | 16% | 0% | 67% | 0% | 0% | 17% | 0% | 0% | 89 |
|  | Punjabi_2 | 0% | 1% | 0% | 70% | 0% | 0% | 29% | 0% | 0% | 103 |
|  | Tamil_0 | 0% | 0% | 0% | 69% | 0% | 0% | 31% | 0% | 0% | 204 |
|  | Telugu_0 | 0% | 0% | 0% | 70% | 0% | 0% | 30% | 0% | 0% | 204 |

We have investigated the difference between inferred haploid parental admixture profiles and diploid admixture profiles of analyzed individuals. For every individual, we have computed an average of parental haploid admixture vectors and calculated the Euclidean distance between the vectors. The average distance for all individuals was 0.092. On average, the largest differences were observed for Toscani, Iberian, Columbian, Mexican Ancestry, and Puerto Rican individuals. In diploid mode, the average value of the Mediterranean component was 60%, South West Asian - 23%, and Northern European - 15%. In haploid mode, the Mediterranean component was 82%, and South-West Asian - 18%. The same situation happened with the Iberian samples. In the diploid mode, the average value of the Mediterranean component was 55%, South West Asian - 19%, and Northern European −24%. In the double haploid mode, the Mediterranean component was 72%, South West Asian - 16%, and Northern European - 12%. Similar inflation of the Mediterranean accompanied by a reduction in the Northern European components.

To investigate the source of this difference, we have identified the largest admixture component for every sample and calculated the difference between the average parental value of this component and its diploid value. The difference ranged from −0.0057 to 0.3063; the mean difference was 0.0684, indicating that haploid Admixture tends to increase the value of the largest component. The magnitude of the increment grows approximately linearly for components <0.87 and then reduces linearly for components above 0.87. Worldwide populations show different trends (Figure 3). For individuals of European and East Asian descent, the trend is positive linear, and the larger the value, the more it is inflated. For South Asian individuals, there is no clear trend. For African and American individuals, there are multiple linear trends. The slope of the line is defined by the relative values of the admixture components. For example, among Columbians, a larger slope corresponds to the individuals where the average ratio of Mediterranean to the sum of Sub-Saharan African and Native American components is 2.07, while a smaller slope corresponds to a 0.56 ratio.

**Figure 3:**
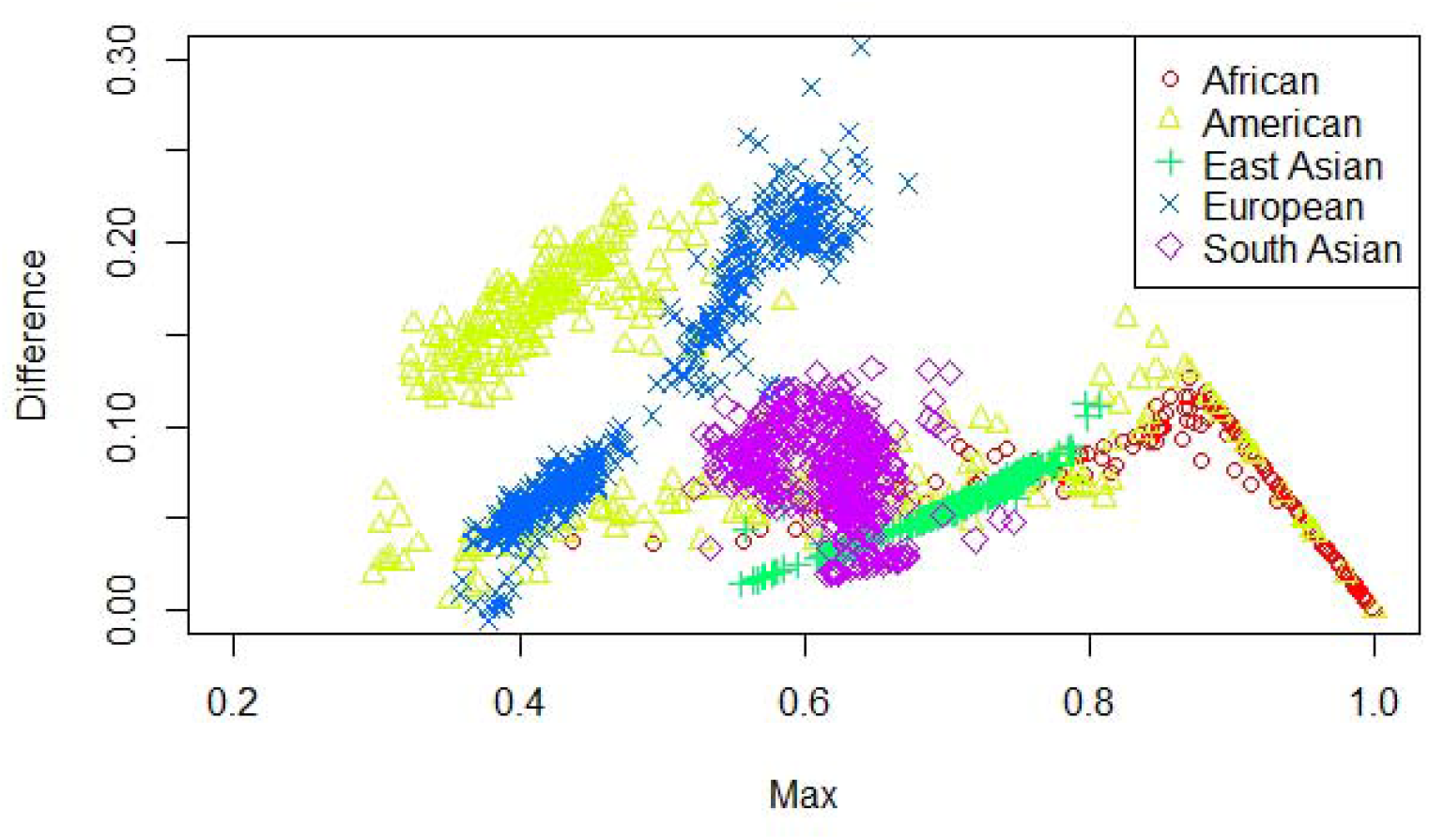
Difference between the largest admixture component in the aggregated haploid and diploid case.

Next, we have investigated the accuracy limitations of the dataset and admixture-based approaches. First, for each individual’s admixture vector, we have calculated the nearest population, based on Euclidean distance between admixture vectors, using the leave-one-out approach; then we have computed fractions of self-hits (the nearest population agrees with the population label of the individual), and fractions of super population self-hits (the agreement is at the level of super-populations). Admixed populations like African American, African Caribbean, Colombian, Mexican in the USA, Peruvian, and Puerto Rican have the lowest assignment accuracy at both population and subpopulation levels. We also noticed that West African populations, such as Yoruba, Esan, Gambian Mandinka, and Mende, are frequently misclassified. Although they speak languages from the same Niger-Congo group, their genomes are not identical [23, 24]. Simplified representation using nine admixture components is artificially “collapsing” those groups to a single point in 9-dimensional space.

**Figure 4:**
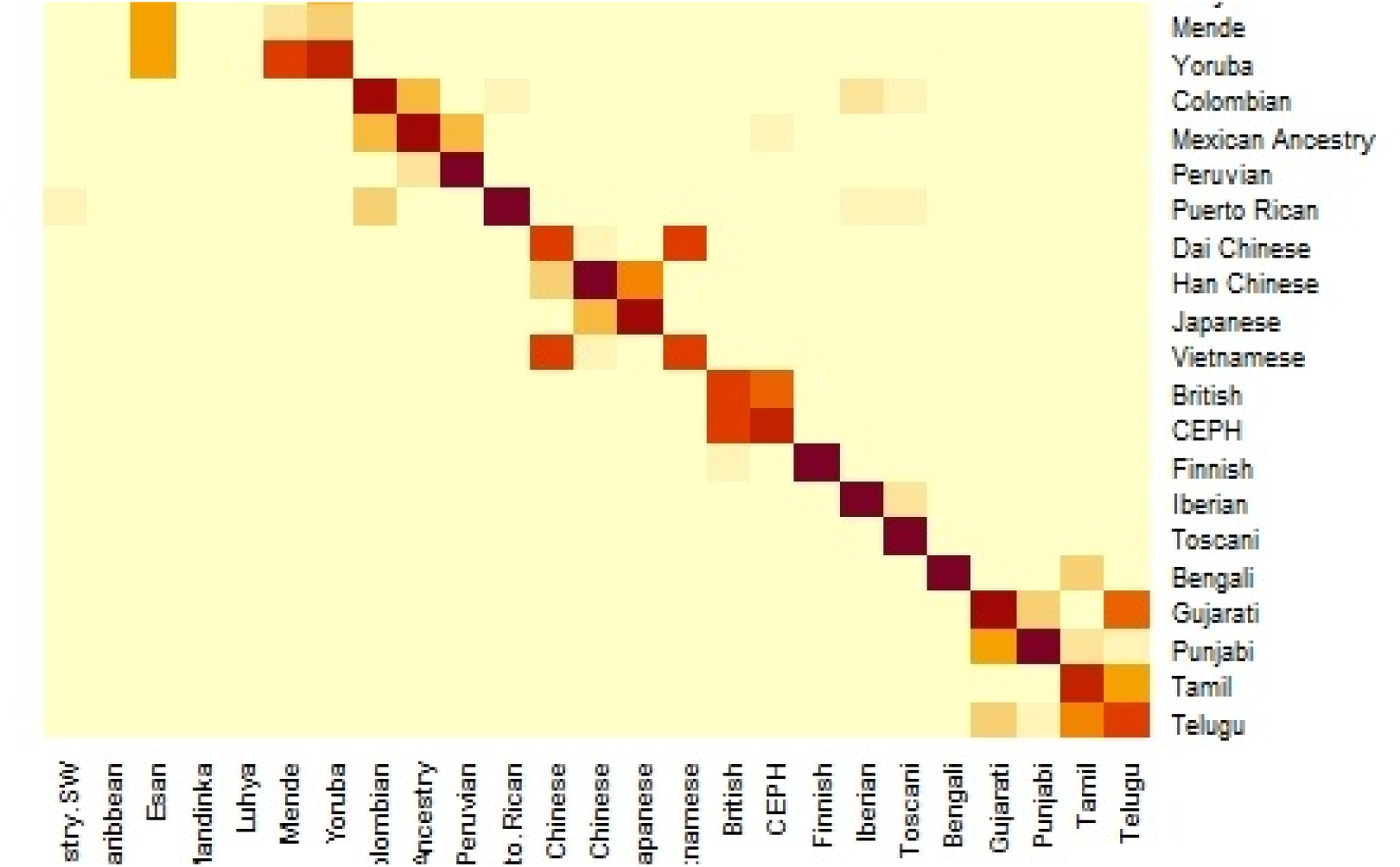
ReAdmix heatmap

GPS and reAdmix (Table 3) analyses suggest that population labels were not 100% accurate but were correct at the superpopulation level. For example, about 30% of Yoruba individuals were mapped to Yoruba reference, but all of them were mapped to one or another African reference. This can be explained by several factors: first, by true heterogeneity of individuals and fuzzy boundaries between populations; second, there may be a human error factor on behalf of the sampled individual or a researcher collecting data; the third factor is the unreported Admixture from different populations. Native and African American individuals appear to be the most affected by these factors; this observation agrees with known historical events. Therefore, we expect to see admixed regions along the genomes of affected individuals.

**Table 3.** GPS and reAdmix

| Super-Population | Population | Number of individuals | GPS, Fraction of populations self-hits | GPS, Fraction of super-population hits | reAdmix, Fraction of populations self-hits | reAdmix, Fraction of super-population hits |
| --- | --- | --- | --- | --- | --- | --- |
| African | African Ancestry SW | 61 | 0.34 | 0.85 | 0.58 | 0.98 |
|  | African Caribbean | 96 | 0.56 | 0.97 | 0.47 | 0.99 |
|  | Esan | 99 | 0.07 | 1.00 | 0.96 | 1.00 |
|  | Gambian Mandinka | 113 | 0.01 | 1.00 | 0.01 | 1.00 |
|  | Luhya | 99 | 0.07 | 1.00 | 0.00 | 1.00 |
|  | Mende | 85 | 0.92 | 1.00 | 0.01 | 1.00 |
|  | Yoruba | 108 | 0.19 | 1.00 | 0.10 | 1.00 |
| American | Colombian | 94 | 0.65 | 0.96 | 0.46 | 0.70 |
|  | Mexican Ancestry | 64 | 0.5 | 0.97 | 0.17 | 0.89 |
|  | Peruvian | 85 | 0.8 | 0.98 | 0.86 | 0.96 |
|  | Puerto Rican | 104 | 0.85 | 0.93 | 0.63 | 0.78 |
| East Asian | Dai Chinese | 93 | 0.62 | 1.00 | 0.62 | 1.00 |
|  | Han Chinese | 208 | 0.71 | 1.00 | 0.25 | 1.00 |
|  | Japanese | 104 | 0.96 | 1.00 | 0.89 | 1.00 |
|  | Vietnamese | 99 | 0.41 | 1.00 | 0.37 | 1.00 |
| European | British | 91 | 0.58 | 1.00 | 0.89 | 0.99 |
|  | CEPH | 99 | 0.57 | 1.00 | 0.13 | 0.99 |
|  | Finnish | 99 | 0.97 | 1.00 | 0.90 | 0.99 |
|  | Iberian | 107 | 0.94 | 1.00 | 0.84 | 0.98 |
|  | Toscani | 107 | 1 | 1.00 | 0.99 | 0.99 |
| Southeast Asian | Bengali | 86 | 0.9 | 1.00 | 0.72 | 1.00 |
|  | Gujarati | 103 | 0.69 | 1.00 | 0.52 | 0.99 |
|  | Punjabi | 96 | 0.59 | 1.00 | 0.46 | 0.98 |
|  | Tamil | 102 | 0.59 | 1.00 | 0.73 | 1.00 |
|  | Telugu | 102 | 0.34 | 1.00 | 0.34 | 1.00 |

**Table 4:**
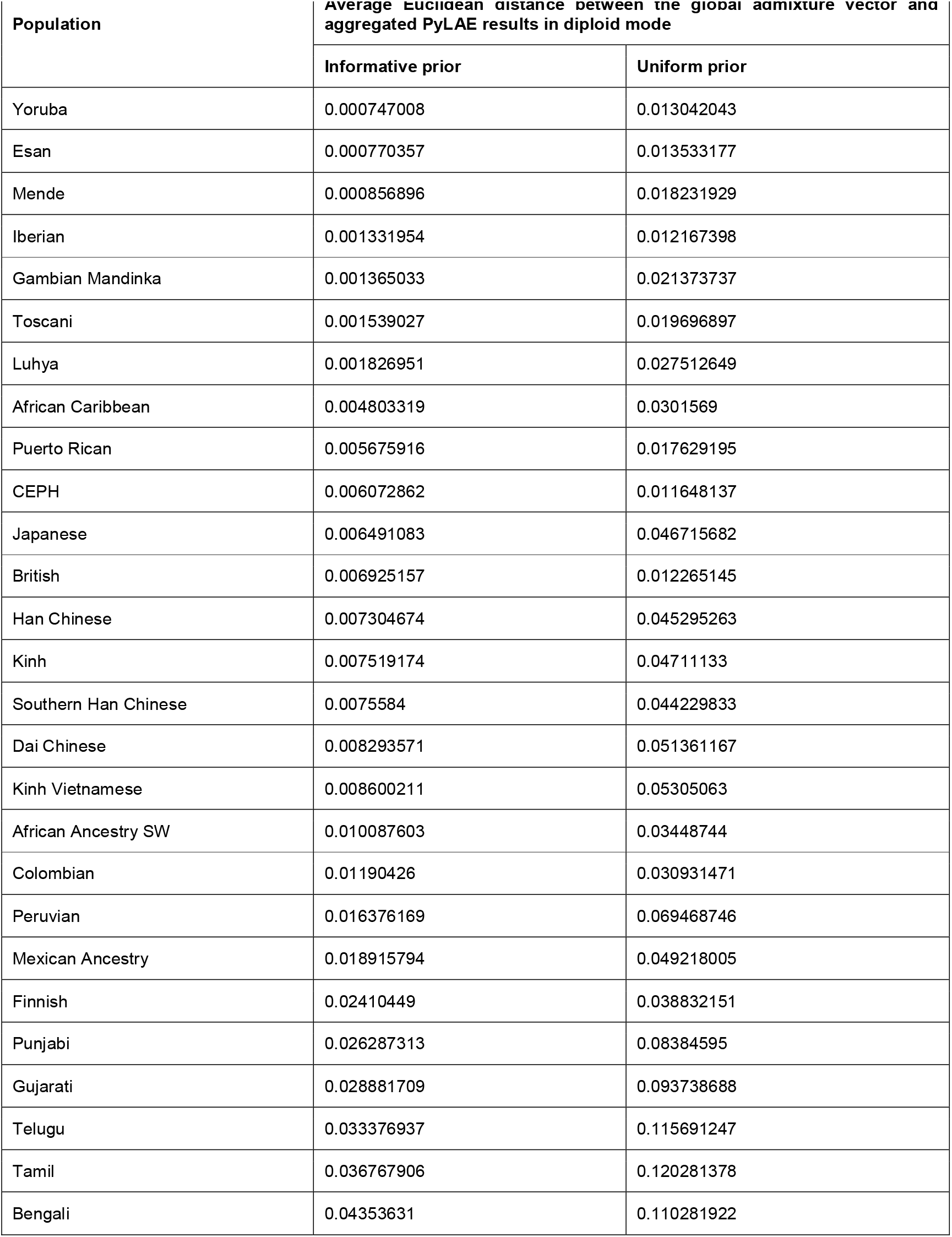
Average Euclidean distance between the global admixture vector and aggregated PyLAE results in diploid mode

**Table 5.**
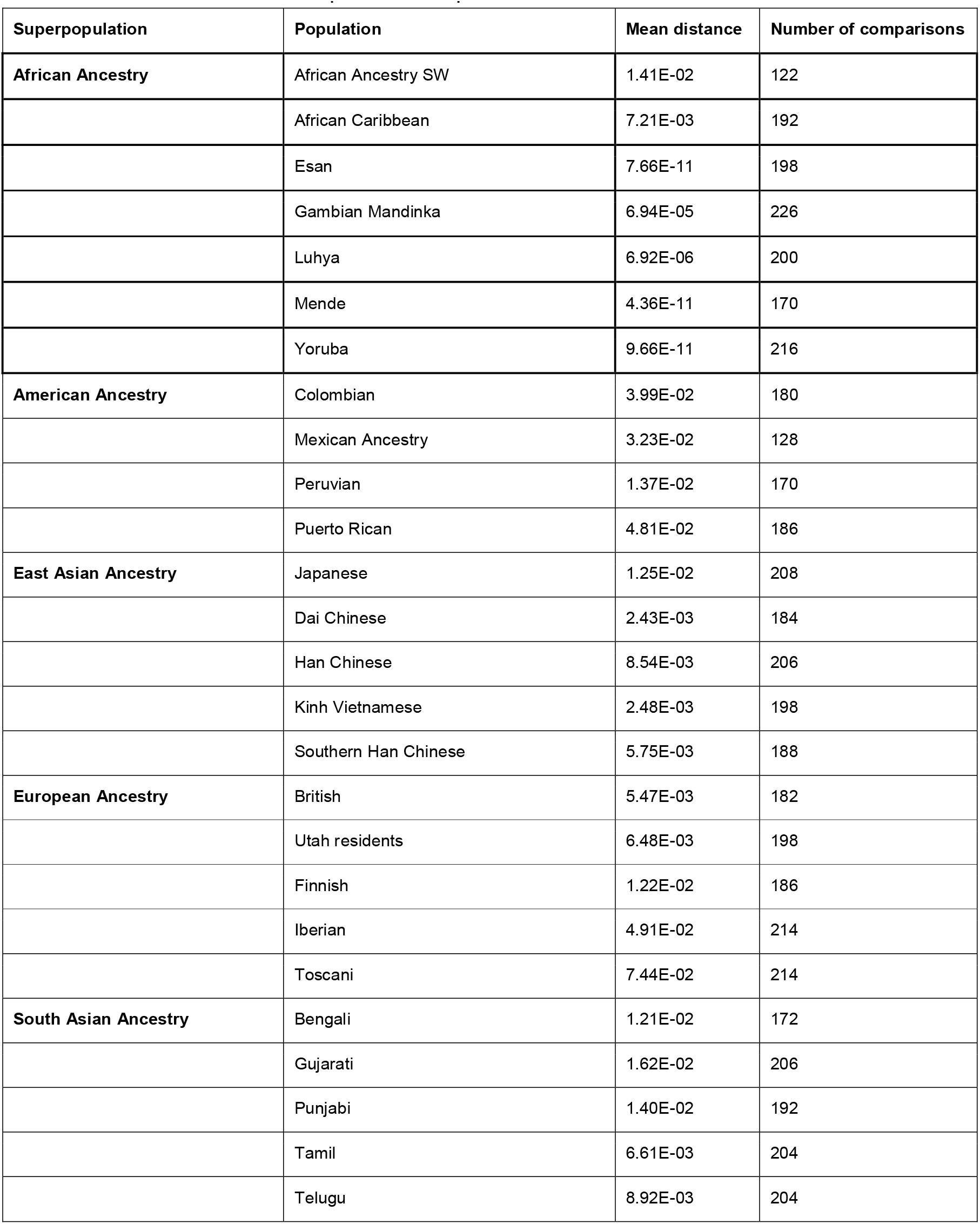
Difference between diploid and haploid Admixture vectors

### 2. Performance of the *PyLAE* algorithm

We have tested the *PyLAE* algorithm on 2504 individuals of the 1kG original dataset using: diploid and haploid versions, with informative and noninformative priors (total four settings). Informative prior is assumed to be equal to the global admixture vector, and the noninformative prior is uniform, assuming equal probabilities for a region to be from any of the K reference populations.

Since we do not know the real locations of regions along the genome, as a formal measure of performance, we chose the agreement with the global admixture vector. For each analyzed individual, we have computed a total fraction of a genome attributed to each of the K components and compared it to the global admixture vector. We have calculated Euclidean distance and Pearson correlation of aggregated results and admixture vectors. The results are shown in Figure 5.

**Figure 5:**
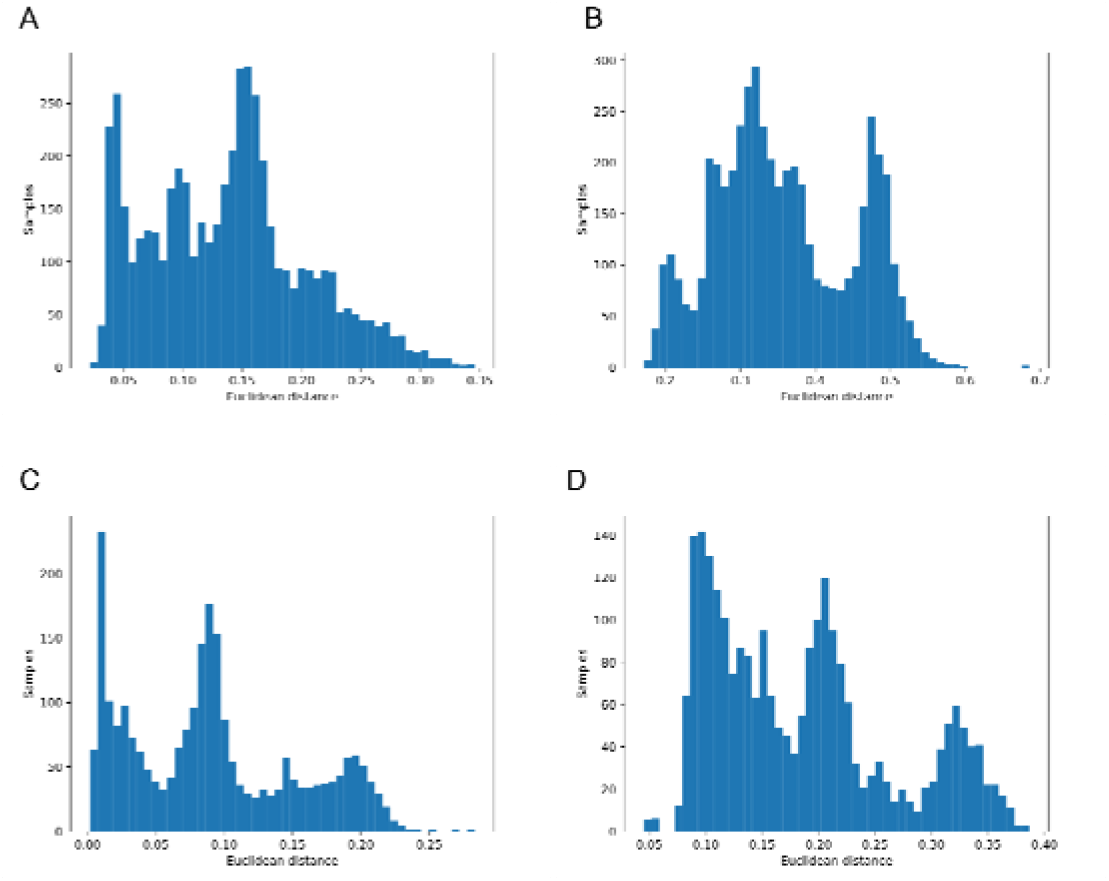
Euclidean distance of aggregated PyLAE results vs admixture vectors for 2,504 from 1000 Genomes. A. ‘Haploid’ mode, with admixture prior B. ‘Haploid’ mode, no admixture prior C. Normal mode, with admixture prior D. Normal mode, no admixture prior.

**Figure 6:**
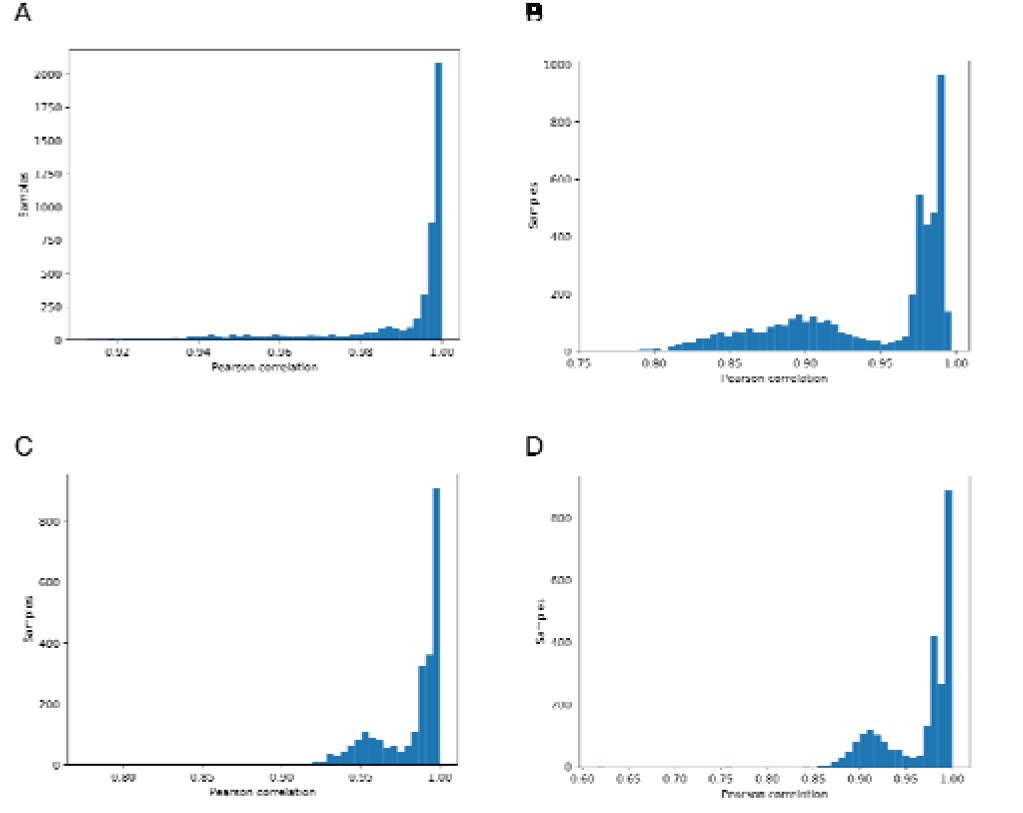
Pearson correlation of aggregated PyLAE results vs admixture vectors for 2,504 from 1000 Genomes. A. ‘Haploid’ mode, with admixture prior B. ‘Haploid’ mode, no admixture prior C. Normal mode, with admixture prior D. Normal mode, no admixture prior.

### 3. Application of Local Ancestry

Using the ANNOVAR [17, 18, 25], and snpEFF [25], we have performed variation annotations and calculated the counts of synonymous and nonsynonymous SNPs. We have conducted this analysis in several modes: (1) comparing British, Finnish, and CEU samples with African American and African Caribbean; (2) comparing British, Finnish, and CEU samples with Africans; (3) comparing British, Finnish, and CEU samples with African samples from Africa, selecting Northern European component in European samples and Sub-Saharan African component in African samples; (4) comparing British, Finnish and CEU samples with African American and African Caribbean, selecting Northern European component in European samples and the Sub-Saharan African component in African samples; (5) comparing British, Finnish and CEU samples with African samples from Africa, selecting the Sub-Saharan African component in African samples; (6) comparing British, Finnish and CEU samples with African Americans and African Caribbean, selecting the Sub-Saharan African component in African samples.

Since we are using nine-component representation, all Africans in our datasets were assigned to the same Sub-Saharan African component, while African Americans have various levels of Admixture from other components. Therefore, local ancestry mode only partitions genomes of European and African American individuals.

On average, PyLAE has higher enrichment scores compared to the whole-genome approach. Comparing the enrichment scores between differentially enriched pathways (p-value<0.01), in 91 cases, PyLAE resulted in higher enrichment scores and 41 - in lower scores for African Americans. If the significance cut-off is lowered, PyLAE results in a higher number of differentially enriched pathways (Figure 7).

**Figure 7:**
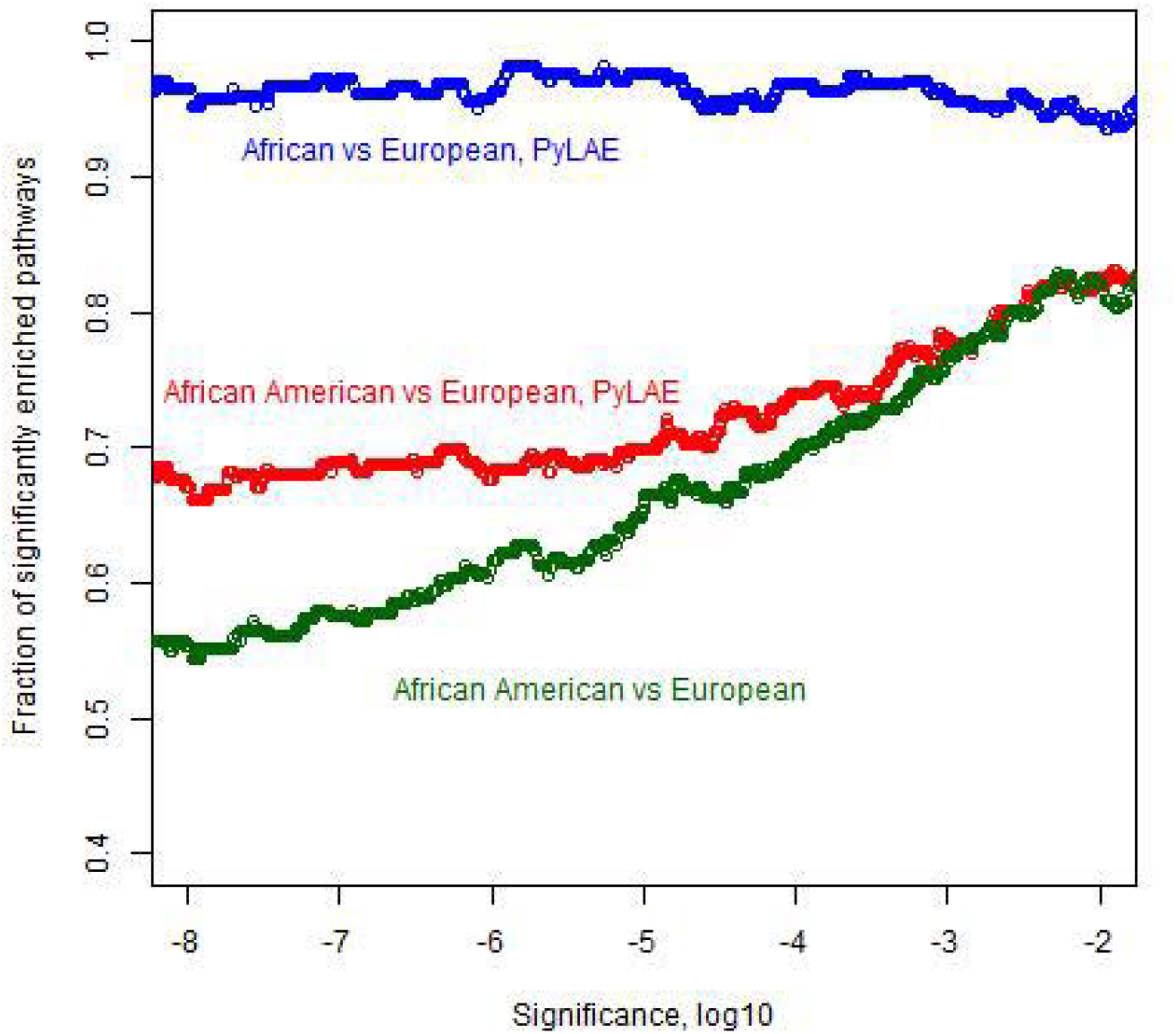
Number of significantly enriched pathways.

Application of PyLAE to African American samples allowed detection of eleven differentially enriched pathways that were not detected when entire genomes of African Americans were used but appear when Europeans are compared with Africans from Africa. According to PyLAE, the following pathways were enriched in nonsynonymous SNPs: **Africans**: Lipoic acid metabolism, Vitamin B6 metabolism, Fatty acid biosynthesis, Notch signaling pathway, Type I diabetes mellitus, Allograft rejection, Cell adhesion molecules (CAMs), Phagosome, Malaria, Viral myocarditis, HTLV-I infection; **Europeans**: Nitrogen metabolism, Mismatch repair, Cell cycle, Glyoxylate and dicarboxylate metabolism, Porphyrin and chlorophyll metabolism, Insulin resistance, Tight junction, Selenocompound metabolism, Oocyte meiosis, Amyotrophic lateral sclerosis (ALS), Protein digestion and absorption.

Dependence on ethnicity efficiency has been reported in processes and conditions related to the above listed pathways: vitamin B6 metabolism [26], fatty acid biosynthesis [27–29], resistance to malaria [30, 31], prevalence of viral myocarditis [32, 33] and amyotrophic lateral sclerosis [34, 35]. According to recent studies [31, 36], individuals with African ancestry have a stronger inflammatory response and reduced intracellular bacterial growth. Maintaining a strong immune response can have adverse side-effects, such as autoimmune disease. Therefore, phagosome efficiency declined after migration out of Africa since Europeans were exposed to fewer pathogens, resulting in reduced immune response over generations.

### Using PyLAE with different genomes and/or sets of markers

A different set of putative ancestral populations or a different set of markers require additional processing. First, we need to collect a database of putatively un-admixed individuals. If there is an existing validated set of ancestry informative features, these markers should run the Admixture in supervised mode. For each self-reported ancestry, samples should be clustered based on their admixture profiles to identify subgroups within each self-reported ancestry. These subgroups are then examined using information about the history of the studied population, and the most representative subset is retained. Then, putative ancestral populations (from 15 to 20 individuals per group) are generated for every ancestry. The validity and stability of the ancestral populations are evaluated using 1) PCA, 2) leave-one-out supervised Admixture, and 3) by application of supervised Admixture to the original dataset.

## Conclusions

*PyLAE* is an easy-to-use and relatively fast tool to perform ancestry decomposition of the sample. As input, it requires a) unadmixed reference split into components of interest, b) prior probabilities obtained from supervised Admixture on the components of interest, or you can use it without prior probabilities (optional, but it will increase the accuracy). *PyLAE* assigns ancestry label to genomic regions. We showed the utility of our approach for the identification of selection signals.

## Availability and requirements

Project name: Python Local Admixture Estimation (PyLAE)

Project home page: https://github.com/smetam/pylae

Operating system(s): Platform independent, Linux is required for preprocessing Programming language: Python, Shell

Other requirements: Python 3.5 and higher; bcftools (preprocessing)

License: Creative Commons

## List of abbreviations

1kG: 1000 Genomes
AIM: Ancestry Informative Markers
GWAS: Genome-Wide Association Study
SNP: Single Nucleotide Polymorphism

